# Aberrant Temporal-spatial Patterns to Sad expressions in Major Depressive Disorders via Hidden Markov Model

**DOI:** 10.1101/2021.03.07.433735

**Authors:** Zhongpeng Dai, Siqi Zhang, Hongliang Zhou, Xinyi Wang, Huan Wang, Zhjjian Yao, Qing Lu

**Affiliations:** School of Biological Sciences & Medical Engineering, Southeast University, Nanjing 210096, China; Child Development and Learning Science, Key Laboratory of Ministry of Education, China; Department of Psychiatry, the Affiliated Brain Hospital of Nanjing Medical University, Nanjing 210029, China; Nanjing Brain Hospital, Medical School of Nanjing University, Nanjing 210093, China

## Abstract

**Background:** The pathological mechanisms of Major depressive disorders (MDD) is associated with over-expressing of negative emotions, and the overall temporal-spatial patterns underlying over-representation in depression still remained to be revealed to date. We hypothesized that the aberrant spatio-temporal attributes of the process of sad expressions relate to MDD and help to detect depression severity.

**Methods:** We enrolled a total of 96 subjects including 47 MDDs and 49 healthy controls (HCs), and recorded their Magnetoencephalography data under a sad expressions recognition task. A hidden Markov model (HMM) was applied to separate the whole neural activity into several brain states, then to characterize the dynamics. To find the disrupted spatial-temporal features, power estimations and fractional occupancy of each state were contrasted between MDDs and HCs.

**Results:** Three states were found over the period of emotional stimuli processing procedure. The visual state was mainly distributed in early stage (0 - 270ms) and the limbic state in middle and later stage (270ms - 600ms) of the task, while the fronto-parietal state remained a steady proportion across the whole period. MDDs activated statistically more in limbic system during limbic state (*p* = 0.0045) and less in frontoparietal control network during fronto-parietal state (*p =* 5.38*10^−5^) relative to HCs. Hamilton-Depression-Rating Scale scores was significantly correlated with the predicted severity value using the state descriptors (*p* = 0.0062, *r* = 0.3933).

**Discussion:** As human brain exhibited varied activation patterns under the negative stimuli, MDDs expressed disrupted temporal-spatial activated patterns across varied stages involving the primary visual perception and emotional contents processing compared to HCs, indicting disordered regulation of brain functions. Furthermore, descriptors built by HMM could be potential biomarkers for identifying the severity of depression disorders.

## Introduction

In major depressive disorder (MDD), patients usually exhibit a mood-congruent processing bias especially toward the processing of negative emotional facial expressions (Stuhrmann, Suslow, & Dannlowski, 2011). This kind of mood-congruent bias makes MDD patients over-express negative information (Dalili, Penton-Voak, Harmer, & Munafo, 2015). This phenomenon was characterized with the mechanisms of stimulus processing and specific over-stimulated biological system (Furey, 2011; Roiser, Elliott, & Sahakian, 2012). The response to negative emotional stimulus in visual areas would be regarded as biomarkers for depression (Furey et al., 2013).

A series of event-related potentials (ERPs) studies have shown that the processing of negative emotional information bias could be a potential phenotype for MDD Harris, & Williams, 2018). During the process of negative stimulus task, several ERPs waveforms were identified to be associated with MDD. Early P1 (100-150ms) reflecting fast perceptual process and evoked unconscious emotions in MDD. Middle N170 (150-200ms) was regarded as indispensable component of emotional processing. The amplitude of late P300 (250-600ms) was suggested to be a biological marker for pathophysiological mechanisms. According to these founds, one can see that the negative bias for MDD was distributed over various temporal stages following the onset of emotional stimuli.

Since the different temporal stages for the process of the negative emotion were closely associated with impaired neural mechanism for MDDs, aberrant spatial patterns underlying different periods deserved further analyses. Based on this, literature reported that the responsiveness of brain regions like amygdala and hippocampus, which were involved in limbic system, was associated with depression severity by utilizing sliding time window algorithm (Bi et al., 2019; Suslow et al., 2010). Previous time frequency analysis in sensor level has found that MDD patients expressed increased activations in occipital lobes when dealing with sad faces (Jiang et al., 2019). Neural activities in prefrontal cortices were hypo-activated in MDD patients under the negative emotional stimulus with dynamic connectivity regression (DCR) algorithm (Bi et al., 2016). Besides, the response of the parietal lobe to sad faces also negatively correlated with disorders severity according to micro-state analysis (Mel’Nikov et al., 2018; Soni, Muthukrishnan, Sood, Kaur, & Sharma, 2018). Lots of evidences have suggested that MDD patients exhibited the disrupted spatiotemporal specificity when dealing with sad facial stimulus, nevertheless, studies revealing the transient dynamics in large-scale brain networks were still rare in relation to the task. Thus, further exploration is required to probe the rich dynamic dysfunction underlying MDD of during the whole period for stimuli processing.

A novel data-driven algorithm Hidden Markov model (HMM) could be utilized to characterize the dynamics of the temporal-spatial brain patterns in network level. Relative to traditional window-based approaches, it could capture transient neural signals under pure negative emotion in milliseconds for each subject, without pre-specifying the window length (Quinn et al., 2018; Vidaurre et al., 2016). Besides, it is convenient for HMMs to generate state-wise mean activation maps in large-scale network via the multivariate observation model, as well as to observe the processing of visual perception, decision making and motor response by the sequential spatiotemporal activation maps. Recent study also showed that impaired brain dynamics could be characterized not only in limited targeted regions but also in the large-scale brain networks via HMMs (Charquero-Ballester et al., 2020). Furthermore, dynamic descriptors such as fractional occupancy inferred from HMMs were found to correlate with symptoms of schizophrenia patients, emerging the potential of the promotion to other psychosis, like MDD (Zhi, Calhoun, Lv, Ma, & Ke, 2018).

In the present study, we aimed at exploring the abnormal spatiotemporal brain patterns in MDD patients under the negative emotional task. To achieve this, an AE-HMM model was applied on Magnetoencephalogram (MEG) data recorded under the stimulus of sad facial recognition task. MEG with high temporal resolution in milliseconds could be utilized to explore neuropsychological mechanisms of fast neural activity for MDD. The processing of chronological neural activity across visual task was characterized by several brain states whose dynamic descriptors are identifiable for MDD. Furthermore, regression analysis was adopted to explore the ability of these dynamic descriptors to indicate the severity of disorders in MDD. Our study might be a promotion to apply HMM to the field of MDD. We provided a new perspective to the evolution process of negative emotional stimulus over the visual task especially in MDD patients.

## Methods

### Participants

One hundred individuals (50 HC and 50 MDD) were enrolled in the Nanjing Brain Hospital between October 2011 and June 2016. All individuals were given Mini international Neuropsychiatric Interview (MINI) to exclude potential MDD from HC. After exclusion in clinical judgment (potential bipolar disorders) and imaging quality (excessive head motion and other artifacts), ninety-six individuals (49 HC and 47 MDD) were recruited in this study. They were all matched in gender, age and education (**Table1**). For MDD, the severity of disease was assessed by professional psychiatrists using Hamilton Depression Rating Scale (HAM-D) and the Diagnostic and Statistical Manual of Mental Disorders, fourth edition (DSM-4). The inclusion criteria were no brain injury, alcohol or drug abuse. All individuals were right-handed and provided with written consent forms. This study was approved by the ethical committee at Nanjing Brain Hospital.

**Table 1.** Demographic and clinical scores of all individuals.

|  | MDD<br>N = 47 | HC<br>N = 49 | *t* | *p*-value |
| --- | --- | --- | --- | --- |
| Gender (male/female) | 24 / 23 | 25 / 24 | 0.094 | 0.844 |
| Age (years) | 31.68 ± 7.54 | 31.53 ± 7.40 | 0.057 | 0.963 |
| Education (years) | 13.57 ± 5.91 | 14.47 ± 4.56 | 0.112 | 0.828 |
| Score of 17-item HAMD | 21.83 ± 5.95 |  |  |  |

Data were presented as the range of mean ± standard deviation (two tailed *t*-tests)

### Stimuli and task

All individuals were engaged in a passively viewing task of emotional faces. Forty sad facial expressions were chosen from the Chinese Facial Expression Video System (Jiang et al., 2019; Jing-Lun, Yao, & Xie, 2007). Each video was presented lasting for 3 seconds. Then a fixation cross picture was also displayed during the rest. An random inter-trial interval of 0.5s, 1s, or 1.5s followed each facial expression.

### Data acquisition and preprocessing

Task MEG data were recorded with a whole-head CTF275 MEG system (VSM Inc) with a 300 Hz sampling rate. Individuals were scanned lying in the supine position in a magnetically shield room. When scanned, individuals were required to adjust head positions if their head motions exceeded 5 mm compared to the initial position. Individuals’ structural T1 images were recorded using a 1.5T GE system and 3D gradient-echo pulse sequence.

MEG data were pre-processed using Matlab-based fieldtrip toolbox (Oostenveld, Fries, Maris, & Schoffelen, 2011). First, Notch filter was used to remove 50Hz power line noise. Then, we excluded trials containing excessive variance with large channel jumps based on visual inspection. No significant difference in the number of excluded trials was found between MDD and HC (MDD:6.1 ± 2.7; HC:5.3 ±3.2). Furthermore, data were decomposed into 275 components using an independent component analysis (ICA) (Jung, Makeig, Westerfield, Townsend, & Sejnowski, 2001). The number of ICA components was equal to the number of MEG sensors and components related to cardiac and muscle artifacts, eye blinks were rejected.

### Source reconstruction

The pre-processed data in sensor space were projected in source space onto a 6mm grid by a Linearly Constrained Minimum Variance (LCMV) beamformer (Woolrich, Hunt, Groves, & Barnes, 2011). The LCMV beamformer normalized grid in MNI space and constructed a realistic head model using participant’s structural MRI. A covariance matrix was calculated across all trials using the spatial filters. Subsequently, the spatial orientations of each epoch were rotated in order to maximize the variance. Source activity was estimated by multiplying spatial filters with sensor-level time series across the whole scale. Subsequently, a multivariate symmetric orthogonalization was applied to attenuate the spatial leakage interferences (Colclough, Brookes, Smith, & Woolrich, 2015).

### Construction of AE-HMM over all subjects

To further explore dynamics in whole brain, MEG data in source space were processed using matlab-embedded HMMAR toolbox (Vidaurre et al., 2016). In this study, an AE-HMM was constructed to exhibit the process of dynamics transition during the emotional face task. Before estimating the model, whole-brain voxel set in source space were projected onto ninety brain regions based on AAL90 template. Subsequently, source time-series were filtered between 1-120 Hz and the amplitude envelope was calculated using Hilbert transition. Data were concatenated across all individuals to constitute a big 3-dimension matrix whose first dimension is the number of individuals, and the second dimension is the number of brain regions, the third dimension is the number of time series. A schematic of the whole process was exhibited in **Figure1**.

**Figure 1.**
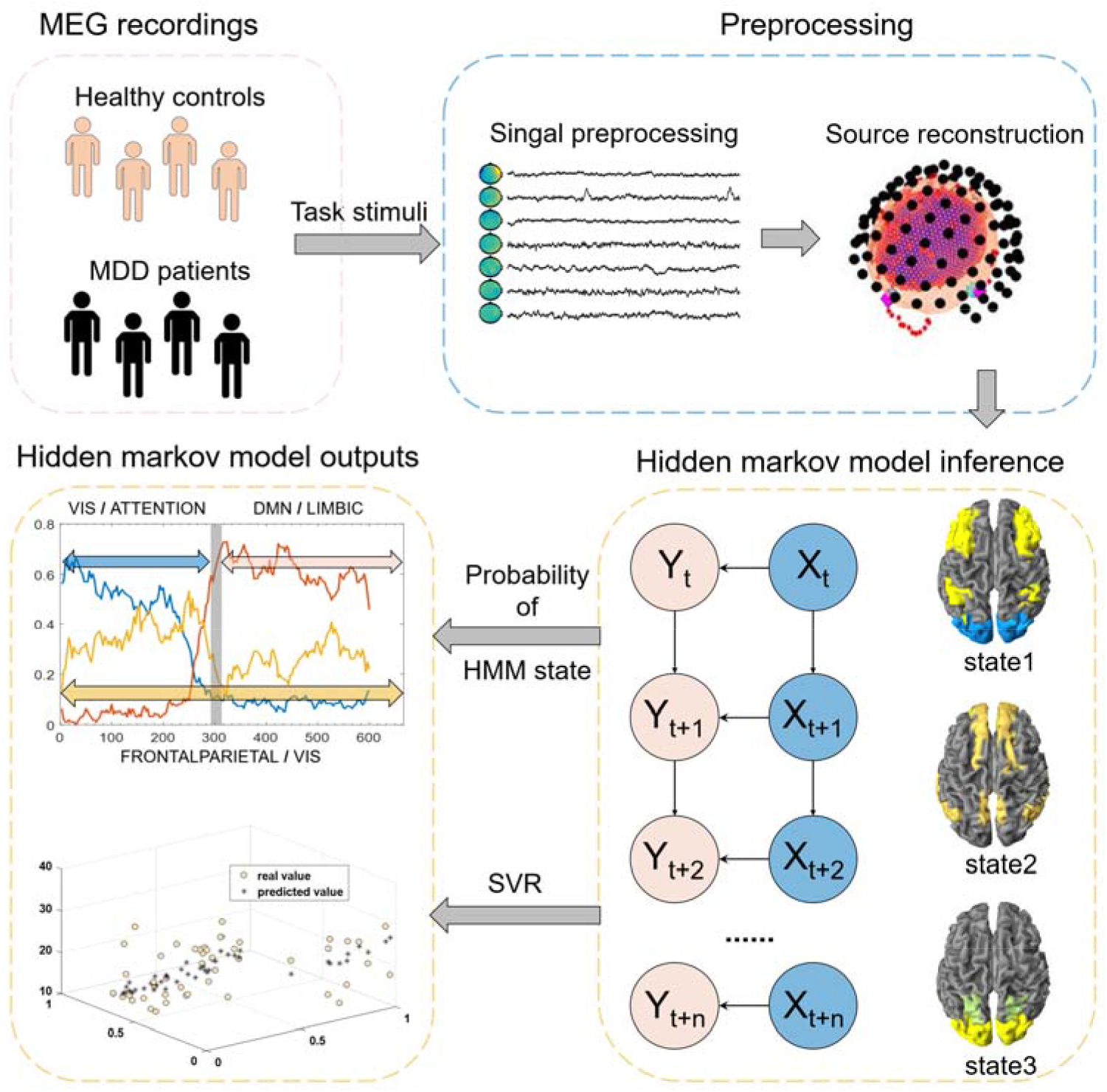
The architecture of the whole pipeline. **(A)** A schematic for the preprocessing of MEG signal applied prior to HMM analysis. **(B)** A schematic for HMM model, in which *X_t_* denotes the brain resides at time point t, while the *Y_t_* denotes the observed data. **(C)** An illustration of brain activation patterns from HMM used to predict disorders severity with Support-Vector-Regression model.

Considering the variance distribution of our data and making sure the lowest free energy, this study denoted K=3 before training the model which also followed findings in Hirschmann’s study (Hirschmann et al., 2020). The observation model subsequently was constructed by training 1000 iterations with pre-defined K states.

### Statistical analysis over state-wise spatial-temporal characters between MDDs and HCs

To characterize the spatial power distribution of each state, covariance matrix was obtained from observation model. The mean activation map was calculated by averaging the prior distributions of the envelope value for brain regions in each state. According to the study in 2019 (Luppi et al., 2019), regions in AAL90 template were divided into seven networks based on dynamic interactions and diversified functions of brain. Each inferred state was shown together with a mean activation for distinct brain networks. Using the time courses of posterior probabilities inferred from HMM, the power activation map of each subject was estimated. Next, non-parametric two-sample *t* tests were utilized over state-wise power activation in network level between MDD and HC groups. The multiple comparisons were controlled via a Bonferroni correction.

In addition, Fractional occupancy (FO) was computed to characterize the dynamics of inferred brain states in each participant. This descriptor is defined as the proportion of each state time spent in the whole time length (Zhi et al., 2018). To further analyze the dysfunction of brain dynamics, FO values in each state were then compared between MDDs and HCs by two-sample independent *t* test.

### Correlational analysis between dynamic characters and MDD severity

To assess the relation between patients’ disease severity and HMM descriptors, a support vector machine regression (SVR) algorithm was applied to regress disease severity in MDD using FO values and HAM-D scores. A Gaussian kernel regression was applied with a leave-one-out-cross-validation (LOOCV). Mean Absolute Error (MAE) was then calculated by average absolute bias between predictive value and real value to assess the predictive performances. Finally, correlational analysis was performed between predicted scores and real HAM-D scores via Pearson correlation.

## Results

### State-wise temporal-spatial patterns for all subjects

In current study, three brain states were identified by AE-HMM over all subjects. As observed in Figure2 A, the fractional occupancy of hidden states at each time point was exhibited through the whole time. Specifically, state1 has a sustained proportion after stimulus onset, lasting about 270ms and characterizing the early stage of task. State2 showed a large occupancy between 270 and 600ms during the middle and latter part of the epoch. State3 exhibited a relatively small but constant occupancy over the whole post-stimulus process.

**Figure 2.**
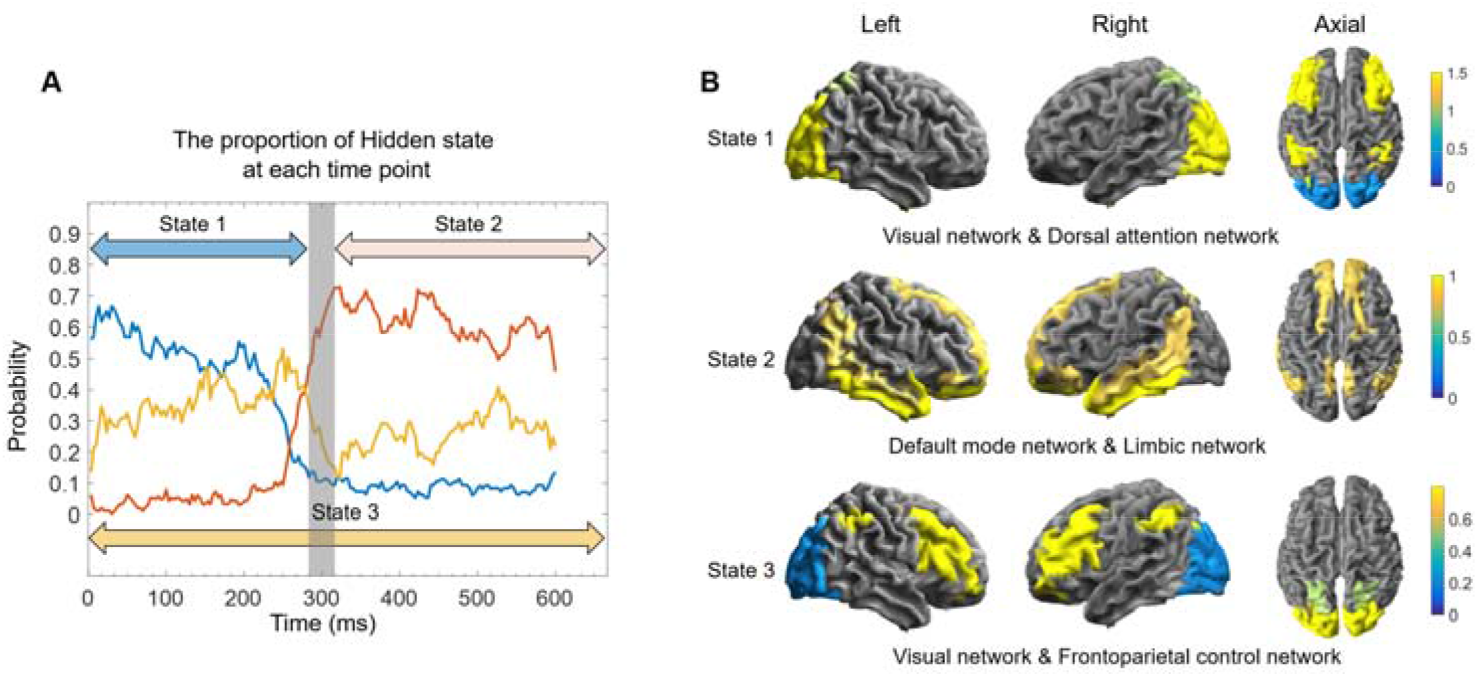
The spatial-temporal brain states of all subjects. **(A)** The averaged probability distribution during the pre-post stimuli across all subjects. The red line denoting state1, pink line denoting state2, yellow line denoting state3. **(B)** The activation pattern characterized by different brain networks in three states. Time series data were normalized to have zero mean, and the regions of top 30% activation level were retained.

In spatial view, as shown in Figure2 B, only brain networks with top 30% activation level was retained to characterize the most active networks of each state. Specifically, state1 showed relatively active neural activation in visual network (VN) and dorsal attention network (DAN), while the brain activity in state2 was mainly occupied by limbic system and default mode network (DMN). State3 was featured with dominant brain activation in VN and fronto-parietal control network (FPN).

### Disrupted spatial-temporal patterns associated with MDD

After comparing network power between MDD and HC groups, network with significant differences were found in two states: limbic system in state2 and FPN in state3. Brain regions in these two networks were illustrated in Figure3 A, relative to HCs, MDD patients manifested significant increased brain activation in limbic system during state2 (*p* = 0.0045, survived after Bonferroni correction) and significant attenuated activity in FPN during state3 (*p* = 5.38*10^−5^, survived after Bonferroni correction) (Figure3 B).

**Figure 3.**
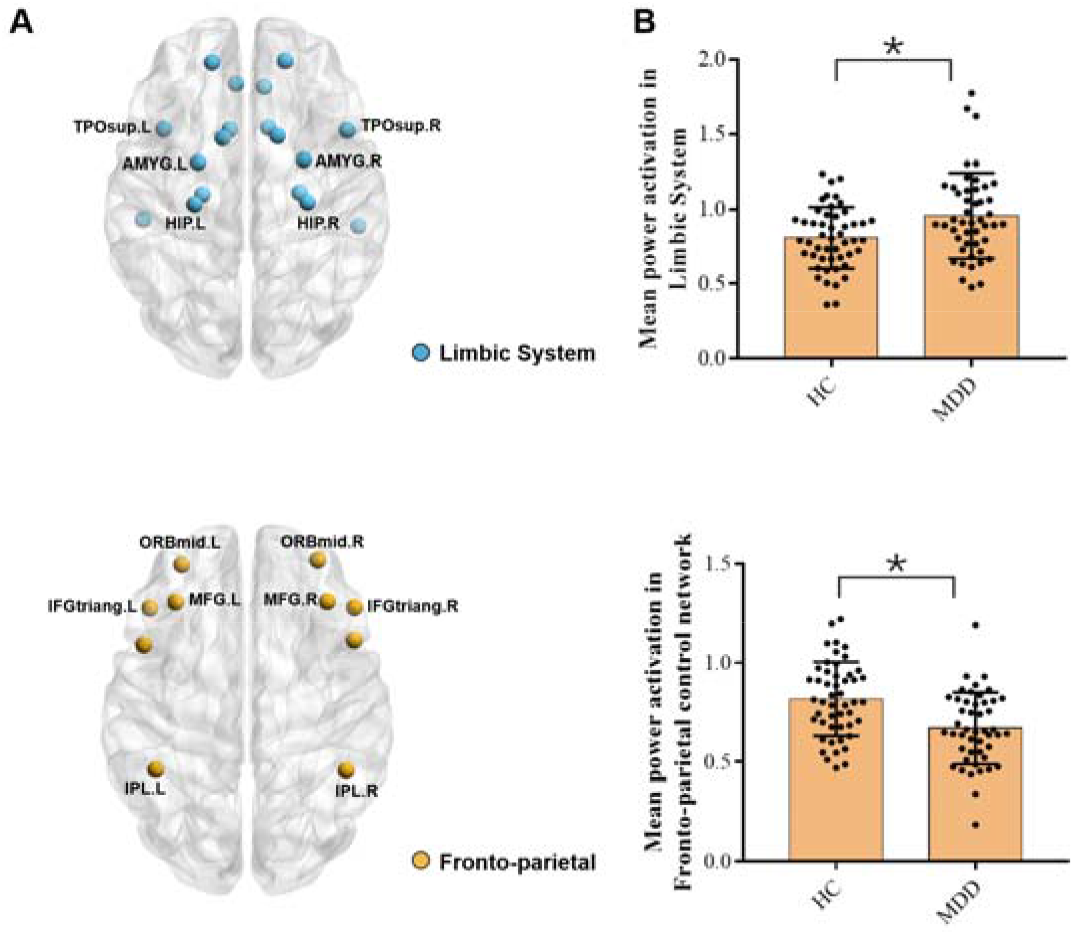
Aberrant spatial patterns for MDD versus HCs. **(A)** Individual network activation maps were compared between MDDs and HCs, and the regions in distinctive networks were shown. The blue nodes represented regions in limbic system and yellow nodes represent regions in FPN (the detailed contents of brain abbreviations in supplementary Table1) **(B)** The distribution of mean network power between HC and MDD groups. The above picture represented mean power activation in limbic system and the below picture represented mean power activation in FPN. \**p* < 0.05 after Bonferroni correction.

For temporal characters, after comparing FO values of each state between HC and MDD groups, MDD subjects exhibited significant higher FO values in state1 (*p* = 0.009, survived after Bonferroni correction) and lower FO values in state 2 versus HCs (*p* = 0.006, survived after Bonferroni correction) (Figure4 A).

**Figure 4.**
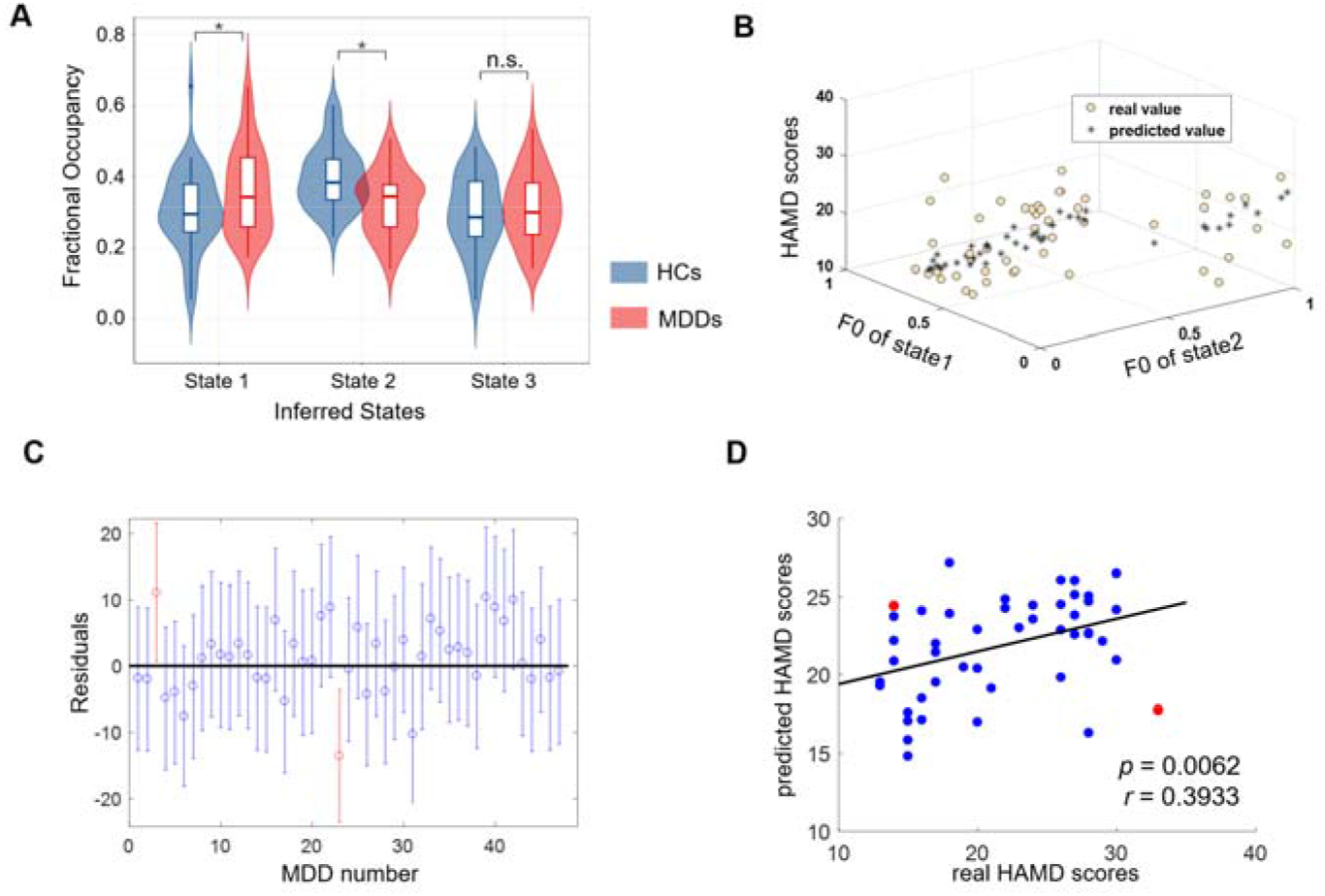
Prediction of disorders severity using FO. **(A)** The violins plot for the distributions of FO values in brain states between HC and MDD groups. \**p* < 0.05 after Bonferroni correction, ns not significant. **(B)** The spatial distribution of clinical scores predicted by SVR model, X axes and Y axes represented FO values of visual state and limbic state respectively, and the magnitude of Z axes expressed the clinical scores, the black hollow dots representing real scores while the green asterisks denoted predicted scores. **(C)** Residuals distributions of predicted scores for each depressed individual, the blue line segment with a middle dot meant the residuals of MAE still in the 95% confidence interval, whereas the red line segment represented the MAE for this individual out of the confidence interval. **(D)** The Pearson correlation with predicted HAMD scores and real scores (*p* = 0.0062, r = 0.3933).

### Dynamic characters related to MDD severity

Since dysfunction of FO was proven to be associated with the depressed disorders, the descriptor was selected as the indicator to the severity of disorders in MDD group. As shown in Figure4 B, the spatial distribution of the real value of HAM-D total scores was consistent with predicted value visually. In statistical, the residuals for each depressed individual were exhibited in Figure4 C, satisfactory predicted outcomes, which meant that the residuals of MAE in the 95% confidence interval, were obtained in 45 subjects out of all the 47 individuals with MDD (Satisfactory ratio: 95.7%). Furthermore, real HAM-D score across all depressed patients were significantly correlated with predicted outcomes in Figure4 D (*p* = 0.0062, r = 0.3933).

## Discussion

This study characterized the aberrant brain activation patterns of mood disorders and found specific state regulating different brain systems through diverse temporal stages of the sad facial recognition task. Besides, dynamic descriptors of FO inferred from each state were regarded as valuable index for individualized severity of depression.

First, the current study revealed the common neural patterns under the negative stimuli for all subjects. We inferred the whole procedure of negative emotions processing into three separate brain states. Specifically, during the early stage of visual stimuli processing (around 0-270ms), subjects showed dominant activation in VN and DAN. During the middle and latter processing stage after the onset of the stimuli (around 270-600ms), subjects manifested more activation in limbic system and DMN. Besides, subjects exhibited a relative small but essential proportion of brain activation in FPN and VN in the whole task (0-600ms). These findings were consistent with what previous literature has reported in visual perception tasks (Quinn et al., 2018). In general, subjects may show an early occipital response after viewing the facial expressions and then gradually transform to the response to the elaborate emotional contents underlying faces by regulating limbic system.

Relative to HCs, MDD patients manifested the aberrant temporal-spatial pattern during the whole process. In early perceptual stage viewing sad expressions, depressed patients manifested statistically decreased FO value in early visual state contrast to HCs. The attenuated FO for early visual perception meant that MDDs spent shorter time capturing negative information from expressions, which suggested the rapid recognition toward sad faces, corresponding to negative emotions biases in previous findings (Almeida, Pajtas, Mahon, Nakayama, & Caramazza, 2013; Sterzer, Hilgenfeldt, Freudenberg, Bermpohl, & Adli, 2011). Besides, in line with previous findings, depressed patients who manifested decreased VN activity might affect visual perception function that was associated with the occurrence and recurrence of mental disorders (Dai & Feng, 2011). Furthermore, previous evidence supported that the DAN often exerts a top-down regulation on the VN to integrate information during the visual perception stage. Thus, this aberrant brain state might represent the dysfunction of top-down mechanism for primary processing, which indicated that MDDs failed to treat negative information comprehensively and their inhibited visual-related cortices made them overexpress negative things and neglect other information (Desseilles et al., 2011; Schupp et al., 2004; Vossel, Geng, & Fink, 2014). In middle and later stage for deep processing of emotional elements. Compared with HC group, depressed patients manifested significant increased FO values as well as hyper-activated power in limbic system of the state. Since limbic-related regions involved amygdala, hippocampus have been considered as the core for processing human expressions configuration and elaborate emotional contents (Almeida et al., 2013; Sterzer et al., 2011), significant higher FO values in limbic state meant that MDDs spent more time indulging in sad contents and over expressing or even ruminating in negative things. Certainly, this kind of aberrant dynamic character underpinned by the impaired pathological neural circuits. Current study reported that statistically elevated activity was observed in limbic system versus HCs during the middle and later stage. Combined with previous findings, we thought that negative information successfully elicited robust responsiveness on limbic system, especially core regions like amygdala (Hall et al., 2014; Suslow et al., 2010). Generally, hyper-activated limbic system in the deeper processing of negative information may be another potential indicator to depressive disorders.

Another important finding was that MDD patients exhibited statistically attenuated power activation of FPN relative to HCs during the whole process. The underlying mechanism underpinning the dysregulated FPN was essential for explaining the neural mechanism of MDD. Previous studies found that FPN was involved in highly adaptive control processes and suggested it could communicate with other systems throughout the brain (Power et al., 2011). In other words, the FPN might regulate other networks to coordinate with specific goals and reduce the goal-disrupting effects brought by other networks. Specifically, evidence supported that both DAN coordinating attention to task stimuli, and limbic system connecting to mental states may be regulated by FPN (Cole, Repovs, & Anticevic, 2014). In present exploratory analyses, although there existed no statistically difference in temporal character between MDDs and HCs, decreased activity of FPN was found in MDDs contrast to HCs. Hypo-activated FPN state may interfere the coordinating function that influences goal-driven brain activities and result in dysregulated brain functional networks. The disrupted spatial patterns of FPN state could also be viewed as a potential pathophysiological biomarker for identifying depression.

Besides, in current exploratory analyses, the FO values of early visual and limbic states were applied as input features of a regression model, which get the predicted outcomes consistent with real HAM-D scores. This finding meant that simple FO parameters describing the aberrant distribution of HMM state occupancies could predict the specific disorder severity for each subject. The satisfactory prediction results of 95.7% may be resulted from the process as we descried above that depressed patients perceive the negative emotion from the stimuli more quickly and then over experience in the sad contents. The overall dysfunction for the distort time allocation implied the defective emotional regulation system underpinned by the impaired neural circuits of MDD. Thus, the dynamic character of FO values could not only discriminate the MDD patients but also indicate the individual depression severity.

In summary, we reconstructed the brain states in the overall process of a passively sad expression recognition task via AE-HMM and found that aberrant temporal-spatial patterns in different process stages like primary visual processing and emotional contents processing were correspond to the different dysfunction of brain functions for MDD respectively. Furthermore, dynamic descriptors of FO values inferred from HMMs could reflect the aberrant dynamism and predict the severity of MDD precisely. Overall, our findings may offer new insights on the pathology of negative emotion process for MDD and predict individual depression severity via a very simple dynamic character.

